# Estrogen Dependent Variation in the Contributions of TRPM4 and TRPM5 to Fat Taste

**DOI:** 10.1101/2025.11.05.686796

**Authors:** Emeline Masterson, Naima S. Dahir, Ashley N. Calder, Yan Liu, Fangjun Lin, Timothy A. Gilbertson

**Author notes:** Corresponding Author: Timothy A. Gilbertson, PhD Professor of Medicine, Department of Internal Medicine UCF College of Medicine, 6850 Lake Nona Blvd Orlando, FL 32827. These authors contributed equally to this research (Emeline Masterson, Naima S. Dahir).

## Abstract

Sex differences in physiology have garnered significant interest of late; however, comparatively little is known about the effects of sex on the function of the peripheral taste system. Previously, we have shown that fat taste functions in a sexually dimorphic manner using molecular, cellular, and behavioral assays, and that a subtype of estrogen receptor (ER) proteins are highly expressed in Type II (receptor) cells. The underlying mechanisms of estrogen’s action, though, remain unknown. Here, we sought to better understand estrogen’s role in fat taste transduction at the molecular level by initially focusing on the transient receptor potential channels types M4 (*Trpm4*) and M5 (*Trpm5*), which we have shown to play roles in estrogen-sensitive fatty acid signaling in taste cells. That is, using *Trpm5*-deficient mice, both males and females in the estrus phase showed significantly reduced FA responses, whereas females in the proestrus phase did not, suggesting that there may be E2-*dependent* TRPM5-*independent* FA signaling in Type II cells. During periods of high levels of circulating estrogen, there was no significant difference in cellular responses to fatty acid (FA) stimuli between *Trpm5*^-/-^ mice and their wild-type counterparts. Moreover, supplemental estradiol enhanced linoleic acid (LA)-induced TRPM5- mediated taste cell activation. Finally, while Type II cells depend on TRPM4 and TRPM5 for FA taste cell activation, proestrus (high estrogen) females showed a greater dependence on a TRPM5-*independent* pathway for fatty acid responsiveness. Together, these results underscore the substantial regulatory role of estrogen in the taste system, particularly for fatty acid signaling. Given that the taste system guides food preferences and intake, these findings may have important implications for understanding sex-specific differences in diet and, ultimately, metabolic health.

**SUMMARY:** The current manuscript shows sex differences in fat taste signaling and identifies functional variations in specific transduction elements in taste cells that respond to sex hormones, such as estrogen, that may mediate differences in peripheral fat taste pathways.

## INTRODUCTION

Males and females have long reported differences in ingestive behaviors and the development of metabolic disorders [1–3]. Women tend to have higher ratios of subcutaneous to visceral fat, as well as a higher prevalence of obesity [4–6]. Despite longstanding research highlighting these sex differences, the exploration of this phenomenon has been limited. [3,7] Investigations into sex-dependent variations in obesity and fat perception have uncovered distinct physiological responses. Specifically, Tanaka et al. (2022) observed a greater reduction in fat taste sensitivity in obese males relative to obese females [8]. Concurrently, Shi et al. (2022) demonstrated that, despite greater weight gain on a high-fat diet, females displayed a lower propensity for hepatic lipid accumulation than males, suggesting potential sex-specific differences in lipid metabolism and partitioning [2]. The consequences of obesity are also sex specific. Recent studies have shown that a high-fat diet exacerbated the cognitive decline in Alzheimer’s disease in females [9]. It is understandable, then, that the taste system, in its role of guiding food intake and dietary preferences, would also display sex differences. In fact, distinct differences in female taste palatability across numerous taste qualities have been demonstrated in both humans and murine models [10–12]. Like many others, we recognize the importance of understanding these sexual disparities in taste and sought to better understand them at the molecular level.

The taste system works as an essential bodily function, recognizing toxins and nutrients that guide our ingestive behaviors. This is accomplished through specialized taste cells that are embedded in our tongues, in taste buds that respond to tastants such as sweet, bitter, umami, salty, sour, and fat. Four distinct taste cell types are differentiated by their responsiveness to different taste stimuli: Type I (glial-like) cells and Type II (taste receptor) cells that respond to sweet, bitter, umami, and fat stimuli. Type III (pre-synaptic) cells contain ion channels for sour taste and may have indirect responses to other tastants, and Type IV (basal/progenitor) cells [13]. The gustatory perception of fat is most commonly experienced through the consumption of dairy products and is exemplified by individual preferences for specific milk types (e.g., whole, 1-2% fat), indicating variations in particular sensitivity to fat stimuli [14]. In the early stages of fat taste, research highlighted the dependence on the palatability of food energy and lipid content [15]. Essential fatty acids (FAs) are both calorically dense and provide a high energy content during typical digestion compared to other proteins and carbohydrates [16]. Since these early studies, the recognition of fat as an essential taste, as the FAs stimulate taste cells, has gained momentum [17]. The rise in metabolic diseases has only added to the push for more research into the recognition, detection, transduction, and transmission of fat taste stimuli in taste cells [18]. Studies have indicated that fat taste stimuli are transduced via a G protein-coupled receptor-mediated pathway that activates second messenger cascades, much like other taste signaling cascades, such as sweet, bitter, and umami [13,19,20].

These investigations have identified several other important signaling molecules, including transient receptor potential melastatin (TRPM) channels, specifically TRPM4 and TRPM5, which are monovalent cation channels [21]. TRPM5 has been well established as a depolarization mechanism in taste cells in response to sweet, bitter, umami, and fat stimuli [22–26]. After its activation by binding Ca^2+^ on two binding sites, a conformational rearrangement opens the ion-conducting pore [27], inducing membrane depolarization leading to stimulation of afferent nerve fibers in the taste system [22–26]. Previous taste research has shown that the ablation of TRPM5 results in a loss of preference for FAs [25,28], less sweet taste preference, and higher glucose tolerance [29,30]. The dynamic responses due to loss or damage of TRP channels have been a topic of exploration for some time.

These TRP channels are expressed in various tissues and have been linked to numerous metabolic diseases [31–33]. Previous studies have shown that other TRP channels exhibit sex differences in non-taste tissues; for example, TRPM2 provided neuroprotection against ischemia in males but not in females [34]. The upregulation of TRPV6 is mediated by estrogen [35]. Cholesterol, the precursor for steroid hormones, has been shown to directly regulate the function of TRPV4, TRPM3, and TRPM8 [35–37]. Female hormones, specifically, have also been shown to influence physiological functions via specific TRP channels; TRPV1 sensitization involves estradiol and estrous cycle-dependent nociception [38]. Progesterone P4 decreases thermosensitivity via TRPV1 [39]. E2 modulates thermoregulation via TRPM8 [40]. TRPM4 expression and function were tightly linked to the estrous cycle in the vomeronasal organ and regulated the relative strength of sensory signals in vomeronasal sensory neurons [41].

The incredible plasticity of the taste system in response to hormonal or dietary changes has been shown in recent studies. Highlighting the roles of hormones, specifically adiponectin and estradiol, in modulating taste receptor sensitivity [11,42]. Additionally, fat taste responsiveness is regulated in part by the sex hormone estradiol [43]. Demographic factors, such as age, sex, and physiological changes, such as obesity, also contribute to variations in taste perception [8,10,44]. This research underscored the plasticity of the taste system in response to sex hormones and highlights the need for further investigation into estrogen’s specific role in the taste system.

Previous studies identified the importance of TRPM5 in the FA-induced taste cell activation and downstream signaling in the Type II taste cells in males [45]. Briefly, long-chain unsaturated FAs interact with G-protein coupled receptor-120 (GPR120) that initiates a signaling cascade beginning with 𝛼𝛼-gustducin to trigger phospholipase C beta 2 (PLC𝛽𝛽2) cleavage of phosphatidylinositol 4,5-bisphosphate (PIP_2_) to inositol 1,4,5-trisphosphate (IP_3_). IP_3_ is then free to stimulate Ca^2+^ release from the endoplasmic reticulum via the IP_3_R3 receptor. The rise in intracellular Ca^2+^ then activates TRPM5, inducing membrane depolarization and pannexin-1 ATP release to stimulate afferent nerve fibers [20,25]. Notably, the downstream fat taste signaling element, TRPM5, was shown to be sensitive to female sex hormones, noting that the administration of E2 or P4 decreased TRPM5 mRNA expression [43]. Though we recently showed similar sex-dependent expression differences in TRPM4 in taste cells [43], the influence of other TRPM channels besides TRPM5 in the fat taste signaling pathway has remained unanswered. Elucidating the precise mechanisms of these receptor-hormone interactions is essential for advancing research aimed at identifying effective interventions for the obesity pandemic [17].

In this study, we highlight sex-specific reliance on the TRPM5-*dependent* FA signaling pathway and estrous cycle-dependent functional roles for TRPM4 and TRPM5. Using ex vivo whole-cell patch clamp recordings and functional calcium imaging in taste cells, we found that TRPM4 and TRPM5 are pivotal in both male and female fat taste signaling. However, their functional activity is synchronized to the female reproductive estrous cycle. Additionally, this cyclical regulation of fat taste via TRPM4 and TRPM5 is dependent upon the ovarian sex hormone, 17β-estradiol. Taken together, the regulation of TRPM4 and TRPM5 function by female hormones in taste cells reveals a previously unknown signaling pathway for studying sex differences in mammalian taste physiology.

## METHOD AND MATERIALS

### Animals

All experiments involving animals were performed in accordance with the American Veterinary Medical Association guidelines and with approval from the Institutional Animal Care and Use Committee at the University of Central Florida. Experiments were performed on male and female estrus-cycled mice. All pups underwent genotyping via standard ear-punch procedures, followed by PCR and electrophoresis to confirm the desired transgenic strain before use. Phospholipase C𝛽𝛽2, a marker for type II taste cells, and Green Fluorescent Protein (PLC𝛽𝛽2-EGFP) transgenic mice were generously provided by Dr. Nirupa Chaudhari (University of Miami School of Medicine). Transient receptor potential melastatin 5 knockout mice (*Trpm5*^-/-^: stock #013068) [45] were purchased from the Jackson Laboratory and then backcrossed with the C57BL/6J strain for at least five generations in our animal facility. Mice were maintained on a 12:12-h light-dark schedule and given *ad libitum* access to chow and water.

### Hormone Assessment

Standard estrus cycling was used to determine relative hormone levels based on cycle stage. To categorize females into high- and low-estrogen groups, those in the early phase of their cycle, during diestrus or proestrus, when estradiol levels are rising and have peaked, were designated the high-E2 group. Those in the later phases of their cycle, estrus or metestrus, when estradiol circulation plateaus, represented the low-E2 group. The standard vaginal lavage was utilized [43]. Briefly, cells from the vaginal canal were collected through plastic Pasteur pipettes prepped with a 0.9% NaCl solution, and the vaginal canal was flushed. The collected cells were used to create a wet smear for light microscopy to determine the cycle stage based on cell number and morphology. To best predict when each mouse would be in a particular phase, vaginal cytology was repeated daily at a consistent time point, and at least ten days before experimentation to ensure typical estrus cyclicality.

### Solution and Reagents

Standard Tyrode’s solution contained (in mM) 140 NaCl, 5 KCl, 1 CaCl_2_, 1 MgCl_2_, 10 HEPES, 10 glucose, and 10 Na pyruvate; pH 7.40 adjusted with NaOH; 305-315 mOsm. The sodium-free Tyrode solution contained (in mM) 280 mannitol, 5 KCl, 1 CaCl_2_, 1 MgCl_2_, 10 HEPES, 10 glucose; pH 7.40 adjusted with TrisOH; 305-315 mOsm. Extracellular solutions in which the calcium concentration was changed (Ca^2+^-free or 0.25 mM CaCl_2_) (in mM) contained 140 NaCl, 5 KCl, 1.75–2 ethylene glycol-bis (β-aminoethyl)-N, N, N′, N′-tetraacetic acid (EGTA), 1 MgCl_2_, 10 HEPES, 10 glucose, and 10 Na pyruvate; pH 7.40 adjusted with NaOH; 305-315 mosM. A potassium-based intracellular solution was used for the measurement of membrane potential and contained (in mM) 140 K gluconate, 1 CaCl_2_, 2 MgCl_2_, 10 HEPES, 11 EGTA, 1.5 ATP, and 0.5 GTP; pH 7.2 adjusted with KOH; 290-300 mOsm. 17β-estradiol (10 nM E2) was purchased from Sigma (St. Louis, MO) and made in 100% ethanol; working concentrations were made from stock on the day of the experiment and dissolved in standard Tyrode’s. Linoleic acid (LA) is a polyunsaturated fatty acid (PUFA) that has been considered the prototypical stimulus for FA taste and representative of other PUFAs [46]. LA was purchased from Sigma (St. Louis, MO) and was prepared as described before.[47] Briefly, LA was prepared as a stock solution (25 mg/ml) in 100% ethanol, stored under nitrogen at ࢤ20 °C, and a working concentration (30 µM) was made immediately before the experiment. LA was dissolved in standard Tyrode’s in all experiments. An antagonist of TRPM5, triphenylphosphine oxide (TPPO; 100 µM), and an antagonist of TRPM4, 9-phenanthrol (9-PHE; 100 μM), were purchased from Sigma (St. Louis, MO) and were made immediately before the experiment. TPPO and 9-PHE were dissolved in dimethyl sulfoxide (DMSO), and the final concentration of DMSO was less than 0.1%. DMSO at the concentrations used here did not affect cell health or cellular responses but was included in the LA-alone solutions.

### Taste Cell Isolation

Individual taste cells were isolated from the tongues of 5 to 12-month-old mice as previously described [19,25,42,48,49]. Briefly, mice were euthanized with CO_2_ exposure and subsequent cervical dislocation, after which their tongues were promptly resected. The tongues were then injected between the muscle and lingual epithelial layers with an enzyme cocktail containing ∼0.2mL of Tyrode’s solution, collagenase A (0.5 mg/ml; Sigma, St. Louis, MO), dispase II (2.0 mg/ml; Sigma, St. Louis, MO) and trypsin inhibitor (1 mg/ml; Sigma, St. Louis, MO). The injected tongues were then incubated in a Tyrode’s bath with bubbled O_2_ for 30 minutes, after which the lingual epithelium was manually removed from the muscle layer and pinned out with the mucosal surface down on a Sylgard™-lined petri dish. This epithelium was then incubated in divalent cation-free (Ca^2+^- and Mg^2+^-free) Tyrode’s solution containing 2 mM 1,2-bis(o-aminophenoxy)ethane-N, N, Nʹ, Nʹ-tetraacetic acid (BAPTA, Sigma, St. Louis, MO), followed by a second incubation of the enzyme cocktail at room temperature. For calcium imaging experiments in which individual taste cells are preferred, the epithelium was incubated in Ca^2+^- and Mg^2+^-free Tyrode’s for 4 minutes, followed by 2 minutes in the enzyme cocktail. For patch clamp experiments in which intact taste buds are preferred, the epithelium was incubated in Ca^2+^ and Mg^2+^-free Tyrode’s for 2 minutes, followed by 1 minute of incubation with the enzyme cocktail. After incubation, taste buds or cells were removed by gentle suction using a ∼100-150 µm fire-polished pipette under a low-magnification dissecting microscope. Isolated cells were plated onto 15 mm glass coverslips coated with Corning® Cell-Tak™ Cell and Tissue Adhesive (Corning, PN 354240) for calcium imaging. Isolated buds were plated onto a Cell-Tak™ coated Superfrost microscope slide fitted with an O-ring fabricated from Sylgard™ 184 elastomer (Dow Chemical).

### Patch Clamp Recording

The prepped slides were continuously perfused with a standard Tyrode solution at room temperature (RT) at a flow rate of 3.5 ml/min. Whole-cell recordings were obtained from taste cells maintained in taste buds at RT. Patch pipettes were pulled on a P-1000 Flaming/Brown micropipette puller (Sutter Instrument) from 1.5-mm outer-diameter filamented borosilicate glass tubing, which were then fire-polished to create resistances ranging from 6 to 10 MΩ. Membrane voltage and current signals were generated using an Axopatch 200B and digitized at 10 kHz using a Digidata 1550B and Clampex software (Molecular Devices). For membrane potential measurements from excited taste cells, the current-clamp mode of the amplifier was used while holding the cell at its zero-current level (i.e., at rest). This mode allows for recording depolarizations or dramatic shifts in the membrane potential to a more positive value, eliciting downstream nerve responses. Whereas the voltage-clamp mode, which was utilized to measure taste cell currents in response to stimuli, allows for the recording of the flow of ions into the cell. Inward currents were recorded at a holding potential of −100 mV to facilitate the absence of voltage-gated conductance in taste cells. By utilizing both modes, we were able to clearly observe the activity of ion channels within these taste cells in response to fat stimuli. A 5-s stimulus (5-10 psi) was applied focally to the entire taste cell from a double-barreled theta-glass pipette (Warner Instruments, Holliston, MA) positioned near the cell and delivered by a PicoSpritzer III (Parker Hannifin Corp.) to obtain responses from the same cell. For single-stimulus applications, a micropipette filled with the appropriate stimulus was connected to the PicoSpritzer III and used to deliver a 5-s stimulus. Responses of taste cells were recorded continuously before, during, and after LA application, controlled by the data acquisition software. Response differences to FAs and other stimuli between single taste cells and taste cells contained in a taste bud showed no significant differences, and data were pooled accordingly.

### Calcium Imaging

Radiometric calcium imaging experiments utilizing Fura2-AM were conducted as previously described [19,42,49]. Briefly, prepared coverslips with isolated cells were loaded with intracellular calcium indicator Fura2-AM (Invitrogen) for an hour of incubation before experimentation. To record changes in [Ca^2+^]I, cells were excited at 340 and 380 nm, and emission was recorded at 510 nm using the InCyt Im2 software (Intracellular Imaging Inc.). To identify the subset of Type II taste cells, the GFP-PLC𝛽𝛽2 transgenic lines were utilized, then cells were excited at 490 nm to excite the EGFP-expressing cells, and emission was recorded at 510 nm. The GFP-expressing cells were marked within the software prior to recording the Fura-2 signal. The F340/F380 ratio of each cell was converted to [Ca^2+^]_i_ based on the calcium calibration kit (Invitrogen). Linoleic acid (LA; 30 µM) and a mixture of LA with TRPM4 and TRPM5 blockers, 9-PHE and TPPO, respectively, in random order, were perfused over taste cells at a flow rate of 4 mL/min, followed by 1 min of 0.1% fatty acid-free BSA solution. After each application, Tyrode’s solution was applied until the calcium signal returned to baseline before the next solution was applied.

### Immunofluorescence Staining

Male and female GFP-PLC𝛽𝛽2 transgenic mice were deeply analyzed with isoflurane before transcardiac perfusion, first with saline, followed by a 4% paraformaldehyde (PFA) solution in 0.1 M phosphate buffer (pH 7.4). After this, the tongues were removed and postfixed in PFA for 2 hours, followed by a 30% sucrose solution in 0.1 M PBS (pH 7.4) overnight at 4 °C. Tongues were then separated into anterior sections containing the fungiform papillae and posterior sections containing the circumvallate papillae, which were embedded in the OCT compound (Tissue-Tek®). Frozen coronal sections were cut at 20 μm using a cryostat and mounted onto Superfrost glass slides. Sections were washed with 0.1 M PBS, and nonspecific binding was blocked for 1 h in a blocking solution (10% serum, 0.3% Triton X-100 in PBS). Primary antibodies, KO validated by the vendor [1:100 anti-TRPM4, Alomone Labs (ACC-044)], were diluted in blocking solution and applied to sections for incubation overnight at room temperature. After rinsing in PBS, the sections were incubated with 1:250 diluted goat-anti-rabbit Alexa Fluor 594 (Invitrogen) for 2 h. After washing with PBS, nuclei were stained with 0.5 mg/mL Hoechst 33258 in 0.1% PBS-Tween® 20 (PBST) for 10 minutes at room temperature. The sections were rinsed 3 times in PBS, and coverslips were mounted with Fluoromount G (Southern Biotech). Negative controls were obtained by omitting the primary antibodies only to assess any background signal. Sections were imaged with a laser scanning confocal microscope (Zeiss, LSM710) equipped with 405, 488, 561, and 633 laser lines. Images were processed and analyzed with ImageJ.

## RESULTS

### TRPM5 is nonessential for LA-induced Type II taste cell activation in early-phase estrus cycle females

Two-way ANOVA with the Tukey method for post hoc multiple comparisons was performed to assess the sex differences as well as the effects of the estrus cycle on TRPM5-mediated FA signaling. It had been anticipated that in cells isolated from male *Trpm5*^-/-^ mice, a significant reduction in response to LA-induced inward current and cellular depolarization would emerge when compared to male wild-type mice (Figure 1A, 1D, and 1E). Similarly, the late-phase females with low levels of circulating estrogen also experienced a reduction in response in the *Trpm*5^-/-^ mice compared to wildtype (Figure 1B, 1D, and 1E). Unexpectedly, we found that in early-phase females, which are exposed to high levels of circulating endogenous estrogen, FA responses were not significantly reduced in the *Trpm5*^-/-^ mice compared to their wild-type counterparts (inward current, F(2,70) 14.2, p<0.0001; cellular depolarization, F(2,74) 9.507, p<0.001).

**Figure 1.**
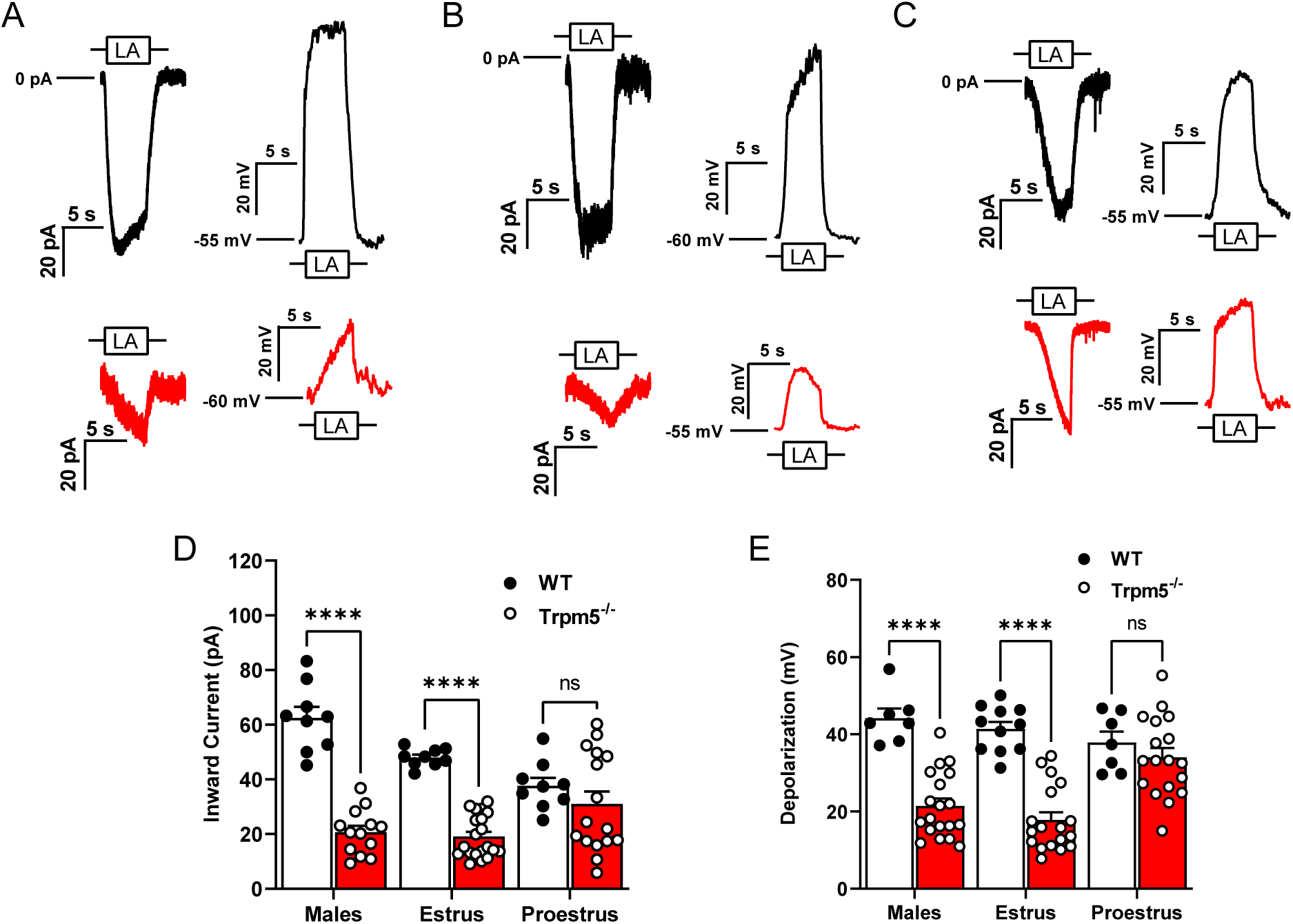
TRPM5 is nonessential for LA-induced Type II taste cell activation in early-phase estrus cycle females. Representative 30 µM linoleic acid (LA)-induced currents and membrane depolarization in wild type (black traces) and *Trpm5^-/-^* mice (red traces) in males **(A)**, females in estrus **(B)**, and females in proestrus **(C)**. Shown are traces of whole-cell *Trpm5^-/-^* currents generated by focal application of LA (30 µM) through a 5-s application via a stimulus pipette. Inward transient currents (at V_h_ = −100 mV) and changes in V_MEM_ (at zero current potential) were recorded from wild-type (PLC𝛽𝛽2-GFP) mice and *Trpm5^-/-^* mice. **(D)** Inward current transients in response to LA in male, female mice in estrus and female mice in proestrus phases of the estrous cycle. Error bars indicate SEM across cells; n = 9 cells for wildtype and n = 13-20 cells from *Trpm5^-/-^* mice. **(E)** Traces depicting total cellular depolarization in response to rapid application of LA (30 µM). Cellular depolarization was recorded from wild-type (PLC𝛽𝛽2-GFP) and *Trpm5^-/-^* mice at resting membrane potential (V_MEM_ = −55 to −60 mV). Error bars indicate SEM across cells; n = 7 – 11 cells from wild type and n = 18 – 19 cells from *Trpm5^-/-^* mice. ****p<0.0001; ns, not significant.

### In Type II cells, estradiol increases LA-induced taste cell activation of the TRPM5-mediated pathway

LA-induced inward currents (Figure 2A) and total cellular depolarization (Figure 2B) were recorded from Type II taste cells in the presence or absence of exogenous E2. A repeated-measures ANOVA with the post hoc Tukey method revealed significant sex- and estrus cycle-dependent differences in taste cell responses to LA. Cells from early-phase females showed no significant difference in LA-induced inward currents (Figure 2A; repeated two-way ANOVA, F(2,18) 18.05, p<0.001) or depolarization (Figure 2B; repeated two-way ANOVA, F(2,18) 34.59, p<0.001). In contrast, Type II cells from both males and late-phase females had significantly increased inward currents and total cellular depolarization in response to LA with added E2 (Figure 2B; V_mem_ 49.57 +/- 3.81). In a manner analogous to the loss of TRPM5, early-phase female responses to LA were insensitive to E2-induced increases in taste cell activation, as seen in males and late-phase females.

**Figure 2.**
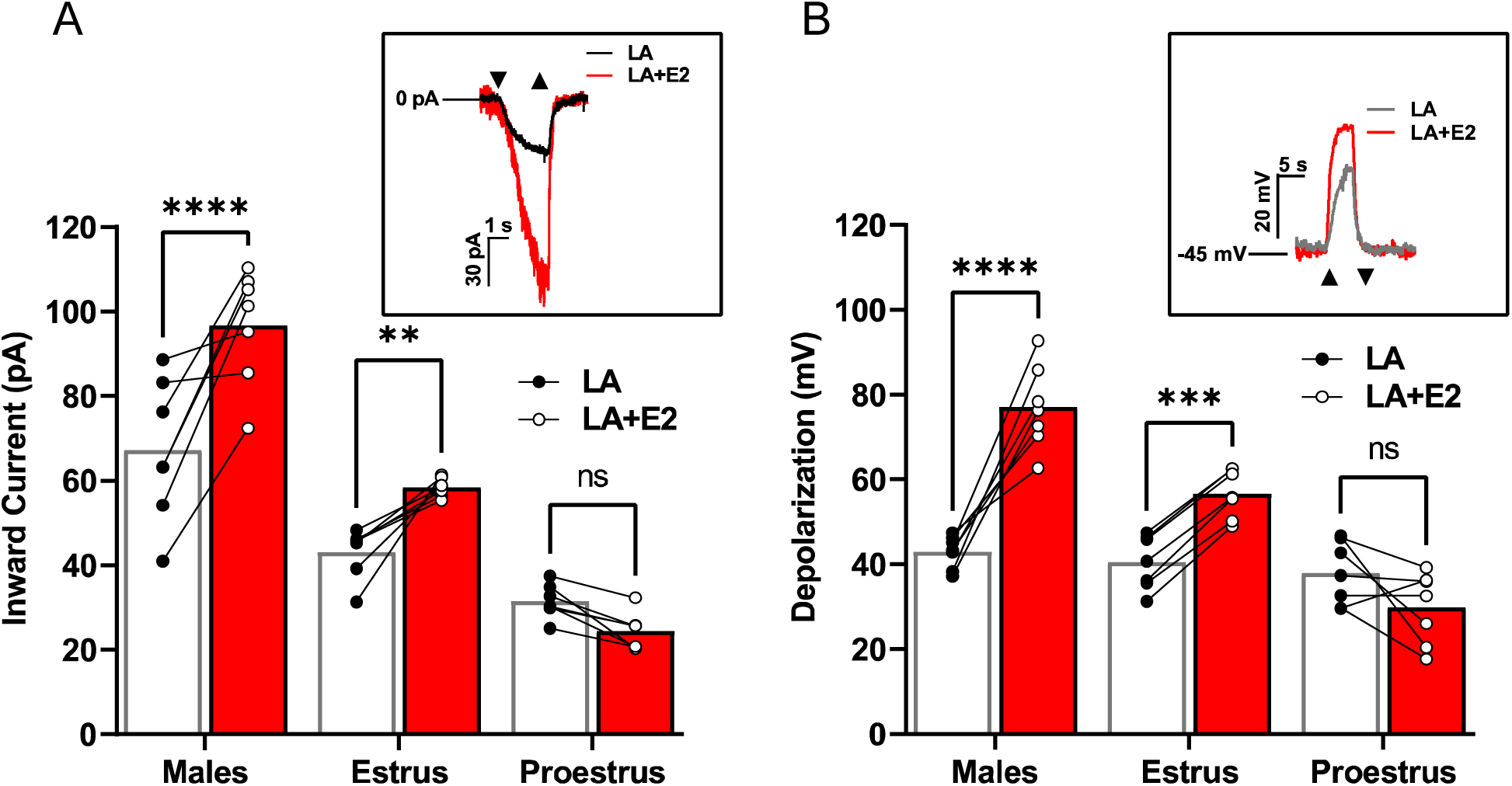
Estradiol increases Type II cell activation in a sex-dependent manner. **(A)** Males and females during the estrus phase of the estrous cycle display an increase in Type II taste cell inward current when FAs were combined with E2 compared to FAs alone. The same cells were used for both stimuli to examine the E2 effect (n = 7 cells for each group). ***Inset*** example inward current trace from a male Type II taste cell that is activated by estradiol with LA + E2 or LA alone. **(B)** Total cellular depolarization is increased in males and females during the estrus phase of the estrous cycle when Type II taste cells are activated by LA with E2 compared to LA alone (holding potential (V_h_) = −100 mV). The same cells were used for both stimuli to examine the E2 effect (n = 7 cells for each group). ***Inset***, example traces of the total cellular depolarization from a male Type II taste cell that is activated with LA with E2 or LA alone (resting membrane potential (V_MEM_) = −45 mV. Cells from females in proestrus showed no enhancement of LA-induced currents or V_MEM_ changes by E2 treatment (n=6 cells per group). ****p<0.0001; ***p<0.001; **, p<0.01; ns, not significant.

### TRPM4 is an essential part of LA-induced fatty acid signaling in Type II cells

To begin, we needed to confirm the colocalization of our PLC𝛽𝛽2-GFP Type II cell marker with TRPM4 proteins via immunofluorescence. We confirmed the presence of TRPM4 protein in the Type II taste cells of male and female mice (Figure 3A). After confirming the presence of TRPM4 in the Type II cells, we sought to confirm its functionality in these LA-mediated pathways in taste cells. To do so, we utilized 9-phenanthrol (9-PHE), a selective TRPM4 antagonist, in males and females and compared the LA-induced Type II taste cell activation. 100 μM of 9-PHE significantly reduced the LA-induced inward currents in Type II cells across all groups (Figure 3B; repeated 2- way ANOVA, F (2,21) 17.86, p<0.0001). Similarly, the total cellular depolarizations from males and late-phase females showed significantly decreased responses to LA, with inhibition of TRPM4 (Figure 3C; repeated 2-way ANOVA, F(2,20) = 30.89, p < 0.0001). Consistent with our previous observations, the early-phase females showed no significant difference in total cellular depolarization response with or without TRPM4 inhibition.

**Figure 3.**
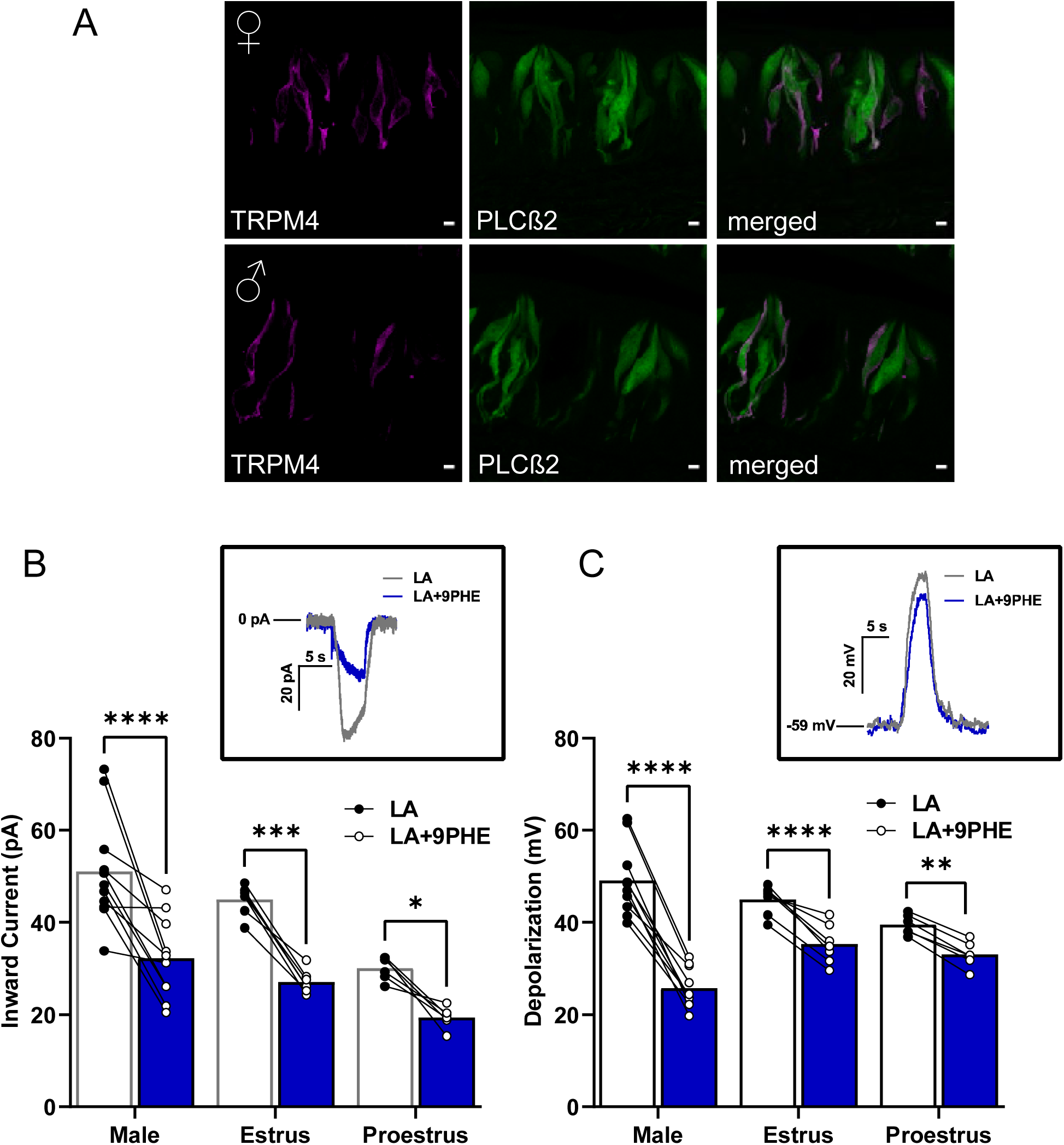
TRPM4 is a key component of fatty acid signaling in Type II cells from male and female mice. **(A)** Immunofluorescence staining from left to right panel showing TRPM4 (magenta), PLC𝛽𝛽2-GFP (green), and merged image. Scale bars represent 5 μm. **(B)** LA-induced inward current in Type II cells is significantly reduced by the TRPM4 channel antagonist 9-phenanthrol (9PHE) in males and females across the estrous cycle. The same cells were rapidly exposed to LA and LA with TRPM4 channel antagonist, 9-phenanthrol (n = 11 cells for males and n = 6 – 7 cells for females). ***Inset*,** inward current trace from male Type II taste cell that is activated by LA or LA with TRPM4 channel inhibitor 100 µM 9-phenanthrol: holding potential (V_h_= −100 mV). **(C)** Total cellular depolarization induced by LA in Type II taste cells is significantly reduced by the TRPM4 channel inhibitor, 9-phenanthrol, in males and females during the estrus phase but not in females during proestrus of the estrous cycle (n = 10 cells for males and n = 6 – 7 cells for females). ***Inset*,** Cellular depolarization of Type II cell rapidly stimulated with LA and LA with the TRPM4 channel inhibitor (100 µM 9-PHE); resting membrane potential. (V_MEM_)= −59 mV).****p<0.0001; ***p<0.001; **p<0.01; *p<0.05.

### Type II taste cells utilize both TRPM4 and TRPM5 for LA-induced taste cell activation

To further evaluate the roles of TRPM4 and TRPM5 in the Type II cell FA signaling pathway, we used selective antagonists, TPPO for TRPM5 and 9-PHE for TRPM4, to inhibit them and assess responses. We performed calcium imaging on single-taste cells isolated from PLC𝛽𝛽2-GFP mice loaded with FURA-2 AM, a radiometric fluorescent dye, and measured the LA-induced intracellular calcium responses. A two-way ANOVA with the Tukey method for post hoc multiple comparisons was performed to compare responses with and without the inhibitors. These results indicated that TRPM4 (9-PHE) and TRPM5 (TPPO) antagonists inhibited LA-induced calcium responses, and the combination of both channel antagonists markedly abolished these responses in both male and female mice (Figures 4A, 4B, 4C). These results indicated that LA-induced calcium activation requires TRPM4 and TRPM5, suggesting an essential role in the inward current response of the taste cells irrespective of sex. As expected, the inhibition of TRPM4 and TRPM5 caused a complete abrogation of calcium responses in the Type II cells. Interestingly, however, the relative effects of the loss of either TRPM4 or TRPM5 varied across the estrus cycle.

**Figure 4.**
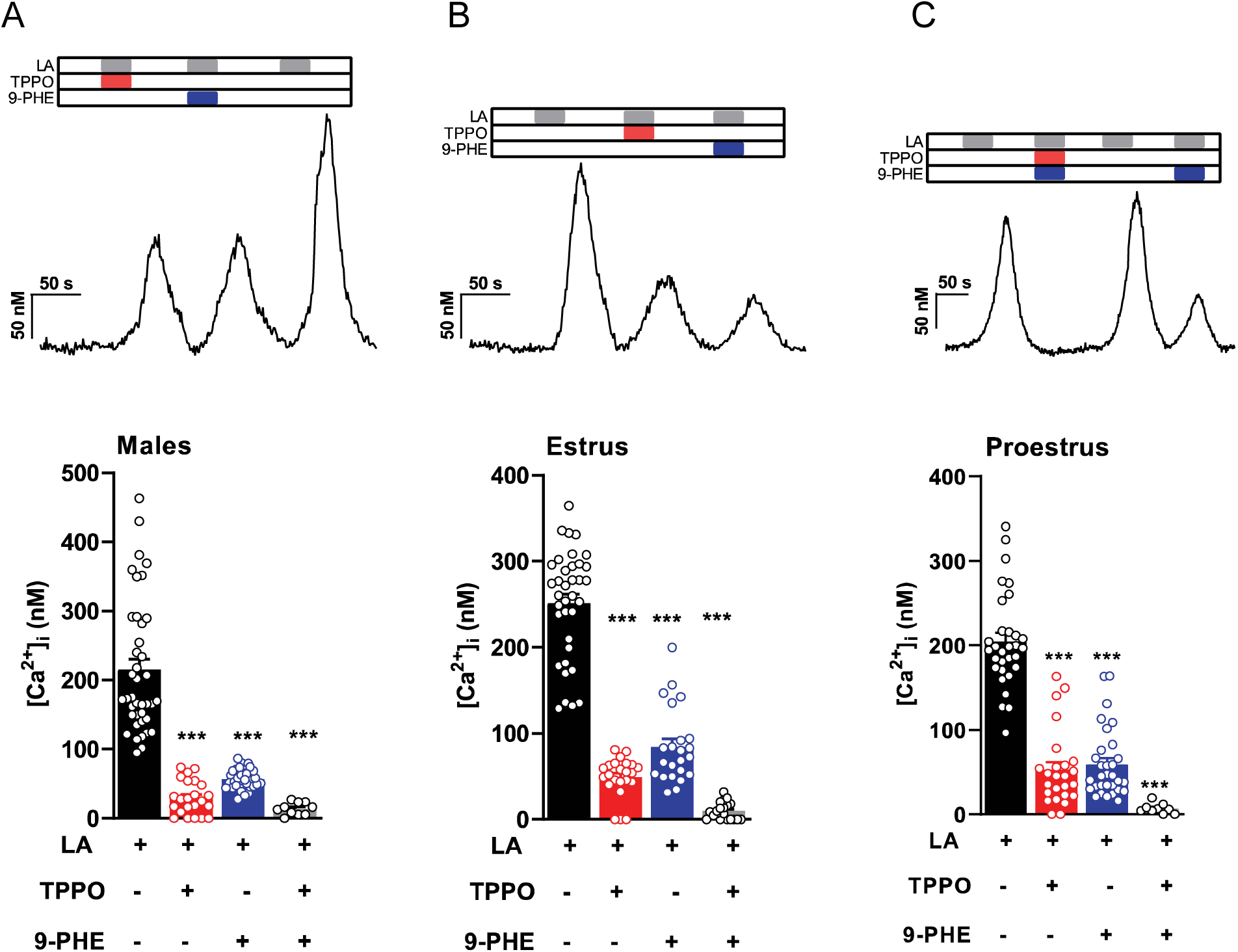
Intracellular calcium rise in Type II taste cells is inhibited by both TRPM4 and TRPM5 antagonists. Representative calcium traces depicting effects of TRPM4 and TRPM5 blockers on LA-induced taste cell responses from Type II taste cells in response to LA, LA with TRPM5 channel inhibitor (100 µM triphenylphosphine oxide [TPPO]), LA with TRPM4 channel inhibitor 100 µM 9-phenanthrol (9PHE), or LA with both TRPM4 and TRPM5 channel inhibitors in males (**A**), females in estrus (**B**), and females in proestrus (**C**). Summary data of intracellular calcium responses induced by LA in the presence or absence of TRPM4 and TRPM5 channel inhibitors are shown from the same group below. TRPM4 and TRPM5 channel inhibitors decreased LA-induced intracellular calcium concentration in Type II taste cells (error bars indicate SEM across cells; n = 23 – 34 cells from males, n = 23 cells from females on estrus, and n = 25 – 31 cells from females on proestrus). *** p<0.001.

### Early-phase females demonstrate enhanced utilization of the TRPM5 independent pathway

To test the ability of early-phase females to utilize a TRPM5-*independent* pathway, we used *Trpm5* knockout mice and the TRPM4 antagonist 9-PHE. We found that the remaining inward current in the *Trpm5^-/-^* males and females in the estrus phase showed no significant difference from the loss of *Trpm5* in the presence or absence of TRPM4 inhibitors. However, the LA-induced inward current from females in the proestrus phase significantly decreased when TRPM4 function was inhibited in addition to the loss of *Trpm5* (Fig. 5A). Similar to the effects in LA-induced inward currents, the LA-induced total cellular depolarization was only affected by TRPM4 inhibition in the females during the proestrus phase (Fig. 5B). These results indicate that LA activation in taste cells shows sex- and estrus-cycle-dependent differences in the role of TRPM4, as well as supporting a functional role for TRPM4 in *Trpm5*^-/-^ mice.

**Figure 5:**
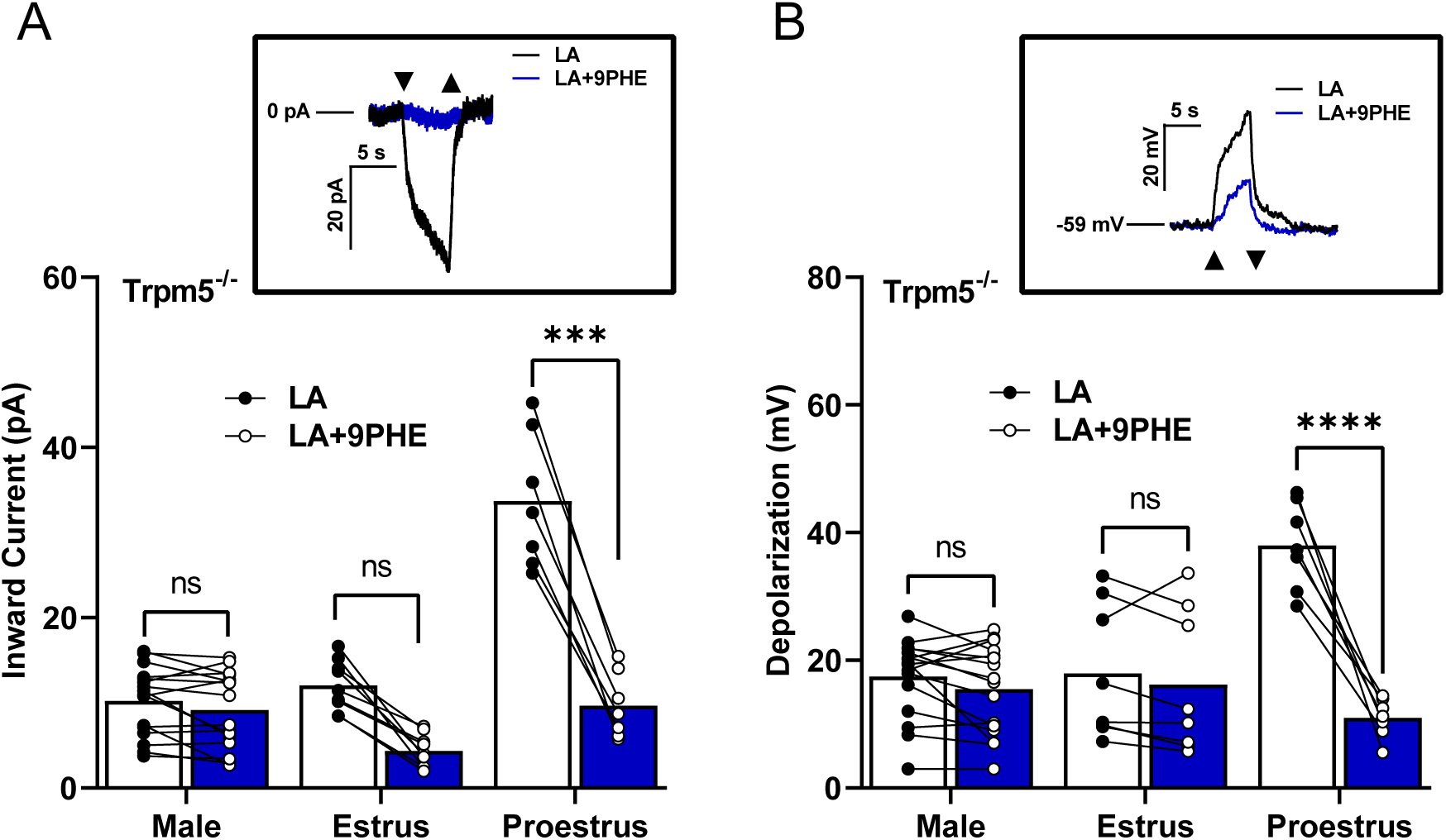
In the absence of normal functioning TRPM5, females in proestrus utilize TRPM4 for LA-induced taste cell activation. LA-induced inward current in female *Trpm5^-/-^* mice during the proestrus phase requires TRPM4. **(A)** Inward current from *T*RPM*5^-/-^*mice stimulated with LA in the presence or absence of TRPM4 channel inhibitor, 100 µM 9-phenanthrol (9PHE; n=16 cells in males, n= 7 – 9 cells in female mice). Inset, an example of inward current traces in a *Trpm5^-/-^*female taste cell during the proestrus phase that is stimulated with LA and LA with 9-phenanthrol, holding potential (V_h_ = −100 mV). **(B)** Total cell depolarization from *Trpm5^-/-^*mice exposed to LA and LA with 9-phenanthrol. In the absence of TRPM5, TRPM4-mediated cellular depolarization is essential in females during proestrus phases (n=16 cells in males, n=7-9 cells in female mice). *Inset*, an example of cell depolarization traces in a *Trpm5^-/-^* female taste cell during the proestrus phase that is stimulated with LA and LA with 9-phenanthrol, resting membrane potential (V_MEM_) = −59 mV. ****p<0.0001; ***p<0.001; ns, not significant.

## DISCUSSION

There have been few, if any, studies that have characterized sex differences in distinct cell types within the peripheral taste system. We provide the initial electrophysiological evidence of sex differences in response to fatty acids. Patch-clamp recordings were made from Type II taste cells, which were unambiguously identified by their intrinsic fluorescence in mice expressing GFP under the control of PLCβ2 [50].

Previous work also demonstrated the significant role that TRPM5 played in the Type II taste cell response to linoleic acid taste perception in male mice [22,25]. Our previous studies examining the effects of sex on the plasticity of fat-taste signaling pathways revealed sex-dependent *Trpm5* expression in taste cells. These differences persisted in females throughout the estrus cycle, with significant fluctuations between early and late estrus phases [49]. In combination, these studies revealed a gap prompting our investigation into the functional role of TRPM5 in mediating sex differences in fat taste. To assess the role of TRPM5, we utilized a knockout model lacking systemic *Trpm5*. In contrast to an earlier study [26], we demonstrated the presence of TRPM4 and IP_3_R3 using PCR, immunohistochemistry, and functional antagonism with a specific pharmacological agent (see Supplementary Figure 1). By combining differential contrast imaging with immunohistochemical staining for TRPM4 and IP_3_R3, we validated their presence in the *Trpm5* knockout mice (Supplementary Figure 1A, 1D). Along with *Trpm5^-/-^* male and normal cycling female *Trpm5^-/-^*mice, we also utilized taste cells from our PLC𝛽𝛽2-GFP transgenic mice, which specifically highlight Type II cells.

### Relative contributions of TRPM4 and TRPM5 vary across the estrous cycle

Similar to previous experiments, we found that Type II cells from male mice required TRPM5 for FA activation (see Figure 1). In electrophysiological experiments, both the TRPM5- mediated inward currents and cellular depolarizations were significantly reduced in male *Trpm5^-/-^*mice. The minor discrepancies between the inward current and depolarization measurements demonstrate the complementary nature of these two techniques. While the inward current measurement quantifies the flow of positively charged ions (such as sodium) into a cell, an inward current initiates cellular depolarization only after a sufficient influx of ions significantly alters the membrane potential. By concurrently measuring cellular depolarization, reflected by a shift in membrane potential towards a more positive value, we can directly observe the relationship between ion flux and cellular activation.

In contrast to males and the majority of earlier experiments, we found that TRPM5 was not solely responsible for taste cell activation in female mice and that TRPM5’s contribution to FA-induced taste cell activation was highly dependent on the estrous cycle. To investigate the role of female cyclic hormones in this TRPM5 FA signaling pathway, these experiments utilized the estrous cycle as a proxy for hormone levels, specifically estradiol (E2). E2 levels exhibit a cyclical pattern, declining during the late estrus phase and rising during the early proestrus phase [51,52]. For example, females in the later estrus phase of their cycle, during which their circulating E2 levels are lower and more similar in magnitude to those of males, showed a significant decrease in both TRPM5-mediated inward currents and cellular depolarizations. However, females in the early proestrus phase showed no significant electrophysiological difference in the FA-induced response between wild-type and *Trpm5* knockout mice. This continued response in the absence of TRPM5 in early-phase females indicates a role for cyclic ovarian hormones in fat taste signaling in female mice. The level of endogenous E2 may influence the use of the TRPM5- mediated fat-signaling pathways. Together, these results support the idea of a TRPM5-*dependent* pathway for fat taste and that this pathway is both sex and estrus cycle-dependent, as females during high endogenous E2 do not require TRPM5 for LA-induced taste cell activation

### In Type II cells, estradiol increases LA-induced taste cell activation of the TRPM5-mediated pathway

Our data suggested a significant role for circulating ovarian hormones in the sex differences seen in taste cell activation. Previous work supported this notion as estradiol increased LA-induced intracellular calcium responses in Type II taste cells [49]. To further explore the role of E2 in the TRPM5-mediated fat-signaling pathway, we assessed the acute effects of E2 in TRPM5-mediated FA taste activation. Previous research had shown that the taste cells did not respond to E2 stimulation alone. However, when applied in combination with FA stimuli (LA+E2), it produced a significant increase in calcium signaling compared to LA alone in taste cells from males and females with low endogenous estrogen [49]. We further examined the interaction between TRPM5-mediated taste cell activation and endogenous E2 levels in females. The experiments showed that the focal application of LA combined with E2 significantly increased both the TRPM5-mediated inward currents and depolarization in wild-type and knockout males, as well as in wild-type and knockout estrus phase females, when compared to LA alone (see Figure 2). Interestingly, there were no significant E2-induced alterations between the wild-type and knockout inward currents and cellular depolarizations observed in females in the proestrus (high E2) phase. Although not significant, the currents in wild-type females during the proestrus phase had a slightly smaller observable magnitude than either the wild-type males or females in the estrus phase. These data indicate that the influence of estradiol on the TRPM5-mediated FA signaling pathway is not only sex-dependent but also estrous-cycle-dependent as well. However, the absence of significant differences in either wild-type or knock-out proestrus females suggests that another mechanism carries the LA-evoked inward movement of Na^+^ ions that promotes taste cell activation.

### TRPM4 is an essential part of LA-induced Type II cell fatty acid signaling

TRPM4 has frequently been proposed to act in concert with TRPM5 to depolarize Type II taste cells in response to sweet, bitter, and umami tastants [13,20,26]. Additionally, our previous work on the *Trpm5* gene expression showed that *Trpm4* expression varied across sexes and the estrus cycle [49]. Given that females during the early proestrus phase of the estrous cycle were able to utilize a TRPM5*-independent* pathway more than their later-phase female counterparts or males, we concluded that the most probable explanation was increased TRPM4 utilization during this phase. Using our genetically identifiable PLC𝛽𝛽2-GFP+ Type II taste cells, we first confirmed the presence of TRPM4 in both males and females (Figure 3A). In patch-clamp experiments, we found that, like other GPCR-mediated tastants, TRPM4 was also an important factor in LA- induced activation of taste cells (Figure 3B, 3C). In both sexes, regardless of the estrous cycle phase, blocking TRPM4 resulted in a significant reduction in LA-evoked inward currents and depolarizations in Type II cells. This confirmed that, as with other tastants, TRPM4 plays an essential role in FA signaling in Type II cells. These data also suggest that early-phase females may be using a TRPM4- and TRPM5-*dependent* pathway. Together, these data show that TRPM4 is an essential part of the FA signaling pathway independent of sex and, most specifically, for the generation of LA-induced inward currents.

### Early-phase females demonstrate enhanced utilization of the TRPM5 independent pathway

To further understand the roles of both TRPM4 and TRPM5 in this FA signaling pathway, we employed ratiometric Ca^2+^ imaging. By perfusing the specific TRPM4 and TRPM5 antagonists, 9-PHE and TPPO, respectively, we directly observed the influence of each channel on LA-induced calcium responses (Figure 4). Inhibiting both TRPM4 and TRPM5 resulted in a nearly complete reduction of the LA-induced intracellular rise in calcium across both sexes and all stages of the estrous cycle. Although not significant, we observed a decrease in the magnitude of the response when the TRPM5 antagonist TPPO is used in both males and late-phase females, compared with the TRPM4 antagonists 9-PHE (Figure 4A, 4B). Early-phase females exhibit similar disruptions in calcium signaling when either TRPM4 or TRPM5 antagonists are applied (Figure 4C). This again suggests an increased reliance on TRPM4-activated pathways in Type II cells in the presence of higher endogenous estrogen.

To further test the role of TRPM4 in this fat-signaling pathway, we combined our *Trpm*5 knockout mice with the TRPM4 antagonist 9-PHE. Interestingly, although trends were evident, males and late-phase females showed no significant difference in LA-induced calcium response magnitudes in the presence or absence of 9-PHE (TRPM4 antagonist) in *Trpm5^-/-^* mice (see Figure 5). However, in Trpm5-deficient mice, early-phase females’ LA-induced calcium responses were greater and highly sensitive to 9-PHE. Thus, TRPM4 appears to play a greater relative role in fatty acid signaling under conditions of high circulating estrogen.

Together, these experiments have demonstrated that the FA signaling, and hence fat taste, is both sex and estrous cycle-dependent, and this dependence is tied to endogenous estrogen (E2) and its role in regulating FA activation in Type II cells. More specifically, in early-phase females, during high levels of endogenous E2, a TRPM5-*independent* pathway via TRPM4 plays a larger role than in males or in females with lower endogenous E2. Consistent with this, we observed a significant decrease in both inward current and cellular depolarization from females in the proestrus phase, suggesting that TRPM4 is the main channel used to activate LA responses in this high estrogen female group. We recently reported in taste cells that *Trpm4* expression is also estrous cycle-dependent, and the highest expression is observed when endogenous E2 is high [49]. Similarly, Eckstein and colleagues [41] found that TRPM4 displayed estrous-dependent function in the vomeronasal organ, another chemosensory organ, and may affect responses to olfactory cues; *Trpm4* is upregulated in cycling females and downregulated when the estrous cycle is abolished by ovariectomy. Estrogen receptors (ERs) are also present in taste cells [49]. Collectively, it is to be expected that TRPM4 significantly contributes to taste-induced activation in Type II cells, particularly in female mice.

## CONCLUSIONS

Sex, a fundamental biological variable, has profound implications for physiology, behavior, and disease susceptibility. Female mammals undergo cyclical hormonal fluctuations that significantly influence their physiological processes, including taste perception. In this study, we explored the roles of TRPM4 and TRPM5 ion channels in mediating sex-specific responses to fat-taste cues. Our findings reveal that both channels play a critical role in fat taste detection in both sexes, but their activity is regulated by the estrous cycle. These results suggest that males and females may employ distinct signaling mechanisms to perceive and respond to dietary fat.

## Data Availability Statement

All relevant data are contained in the manuscript. Acknowledgments: This research was supported by the National Institute of Health award R01DC013318 (tag).

## REFERENCES

1. Mauvais-Jarvis, F. Sex differences in energy metabolism: natural selection, mechanisms and consequences. Nature Reviews Nephrology 2023 20:1 2023–11–03, 20, doi:10.1038/s41581-023-00781-2.

2. Shi, Y.; Wei, L.; Xing, L.; Wu, S.; Yue, F.; Xia, K.; Zhang, D.; Shi, Y.; Wei, L.; Xing, L., et al. Sex Difference is a Determinant of Gut Microbes and Their Metabolites SCFAs/MCFAs in High Fat Diet Fed Rats. Current Microbiology 2022 79:*11* 2022**–**10–08, 79, doi:10.1007/s00284-022-03025-x.

3. Lovejoy, J.C.; Sainsbury, A. Sex differences in obesity and the regulation of energy homeostasis. Obesity Reviews 2009/03/01, 10, doi:10.1111/j.1467-789X.2008.00529.x.

4. Kyrgiafini, M.-A.; Sarafidou, T.; Giannoulis, T.; Chatziparasidou, A.; Christoforidis, N.; Mamuris, Z.; Kyrgiafini, M.-A.; Sarafidou, T.; Giannoulis, T.; Chatziparasidou, A., et al. Gene-by-Sex Interactions: Genome-Wide Association Study Reveals Five SNPs Associated with Obesity and Overweight in a Male Population. Genes 2023*, Vol.* 14, Page 799 2023–03–26, 14, doi:10.3390/genes14040799.

5. Heo, J.-W.; Kim, S.-E.; Sung, M.-K.; Heo, J.-W.; Kim, S.-E.; Sung, M.-K. Sex Differences in the Incidence of Obesity-Related Gastrointestinal Cancer. International Journal of Molecular Sciences 2021*, Vol.* 22, Page 1253 2021–01–27, 22, doi:10.3390/ijms22031253.

6. Fuente-Martín, E.; Argente-Arizón, P.; Ros, P.; Argente, J.; Chowen, J.A. Sex differences in adipose tissue. Adipocyte 2013–7–24, 10.4161/adip.24075, doi:10.4161/adip.24075.

7. Correa-De-Araujo, R. Serious Gaps: How the Lack of Sex/Gender-Based Research Impairs Health. Journal of Women’s Health 2006, 15, 1116–1122, doi:10.1089/jwh.2006.15.1116.

8. Tanaka, A.; Mochizuki, T.; Ishibashi, T.; Akamizu, T.; Matsuoka, T.-a.; Nishi, M. Reduced Fat Taste Sensitivity in Obese Japanese Patients and Its Recovery after a Short-Term Weight Loss Program. Journal of Nutritional Science and Vitaminology 2022/12/31, 68, doi:10.3177/jnsv.68.504.

9. Gannon, O.J.; Robison, L.S.; Salinero, A.E.; Abi-Ghanem, C.; Mansour, F.M.; Kelly, R.D.; Tyagi, A.; Brawley, R.R.; Ogg, J.D.; Zuloaga, K.L., et al. High-fat diet exacerbates cognitive decline in mouse models of Alzheimer’s disease and mixed dementia in a sex-dependent manner. Journal of Neuroinflammation 2022 19:*1* 2022**–**05–14, 19, doi:10.1186/s12974-022-02466-2.

10. Barragán, R.; Coltell, O.; Portolés, O.; Asensio, E.M.; Sorlí, J.V.; Ortega-Azorín, C.; González, J.I.; Sáiz, C.; Fernández-Carrión, R.; Ordovas, J.M., et al. Bitter, Sweet, Salty, Sour and Umami Taste Perception Decreases with Age: Sex-Specific Analysis, Modulation by Genetic Variants and Taste-Preference Associations in 18 to 80 Year-Old Subjects. Nutrients 2018, 10, 1539, doi:10.3390/nu10101539.

11. Stone, B.T.; Rahamim, O.M.; Katz, D.B.; Lin, J.-Y. Changes in taste palatability across the estrous cycle are modulated by hypothalamic estradiol signaling. bioRxiv 2024–04–02, 10.1101/2024.04.01.587593, doi:10.1101/2024.04.01.587593.

12. Skoczek-Rubińska, A.; Chmurzynska, A.; Muzsik-Kazimierska, A.; Bajerska, J.; Skoczek-Rubińska, A.; Chmurzynska, A.; Muzsik-Kazimierska, A.; Bajerska, J. The Association between Fat Taste Sensitivity, Eating Habits, and Metabolic Health in Menopausal Women. Nutrients 2021*, Vol.* 13, Page 4506 2021–12–16, 13, doi:10.3390/nu13124506.

13. Roper, S.D.; Chaudhari, N.; Roper, S.D.; Chaudhari, N. Taste buds: cells, signals and synapses. Nature Reviews Neuroscience 2017 18:*8* 2017**–**06–29, 18, doi:10.1038/nrn.2017.68.

14. Andersen, C.A.; Nielsen, L.; Møller, S.; Kidmose, P. Cortical Response to Fat Taste. Chemical Senses 2020/05/21, 45, doi:10.1093/chemse/bjaa019.

15. A, D. Taste preferences and food intake - PubMed. Annual review of nutrition 1997, 17, doi:10.1146/annurev.nutr.17.1.237.

16. Kim, M.-A.; Kim, S.-M.; Lee, H.-S.; Kim, M.-A.; Kim, S.-M.; Lee, H.-S. Oral/taste sensitivity to non-esterified long-chain fatty acids with varying degrees of unsaturation. Food Science and Biotechnology 2023 33:*3* 2023**–**12–19, 33, doi:10.1007/s10068-023-01502-y.

17. Şeref, B.; Yıldıran, H. A new perspective on obesity: perception of fat taste and its relationship with obesity. Nutrition Reviews 2025/02/01, 83, doi:10.1093/nutrit/nuae028.

18. Khan, A.S.; Keast, R.; Khan, N.A. Preference for dietary fat: From detection to disease. Progress in Lipid Research 2020/04/01, 78, doi:10.1016/j.plipres.2020.101032.

19. Liu, Y.; Xu, H.; Dahir, N.; Calder, A.; Lin, F.; Gilbertson, T.A. GPR84 Is Essential for the Taste of Medium Chain Saturated Fatty Acids. Journal of Neuroscience 2021–06–16, 41, doi:10.1523/JNEUROSCI.2530-20.2021.

20. Gilbertson, T.A.; Khan, N.A. Cell signaling mechanisms of oro-gustatory detection of dietary fat: Advances and challenges. Progress in Lipid Research 2014/01/01, 53, doi:10.1016/j.plipres.2013.11.001.

21. Ullrich, N.D.; Voets, T.; Prenen, J.; Vennekens, R.; Talavera, K.; Droogmans, G.; Nilius, B. Comparison of functional properties of the Ca2+-activated cation channels TRPM4 and TRPM5 from mice. Cell Calcium 2005/03/01, 37, doi:10.1016/j.ceca.2004.11.001.

22. Pérez, C.A.; Huang, L.; Rong, M.; Kozak, J.A.; Preuss, A.K.; Zhang, H.; Max, M.; Margolskee, R.F.; Pérez, C.A.; Huang, L., et al. A transient receptor potential channel expressed in taste receptor cells. Nature Neuroscience 2002 5:*11* 2002**–**10–07, 5, doi:10.1038/nn952.

23. Damak, S.; Rong, M.; Yasumatsu, K.; Kokrashvili, Z.; Pérez, C.A.; Shigemura, N.; Yoshida, R.; Mosinger, B.; Glendinning, J.I.; Ninomiya, Y., et al. Trpm5 Null Mice Respond to Bitter, Sweet, and Umami Compounds. Chemical Senses 2006, 31, 253–264, doi:10.1093/chemse/bjj027.

24. Zhang, Z.; Zhao, Z.; Margolskee, R.; Liman, E. The Transduction Channel TRPM5 Is Gated by Intracellular Calcium in Taste Cells. Journal of Neuroscience 2007–05–23, 27, doi:10.1523/JNEUROSCI.4973-06.2007.

25. Liu, P.; Shah, B.P.; Croasdell, S.; Gilbertson, T.A. Transient Receptor Potential Channel Type M5 Is Essential for Fat Taste. Journal of Neuroscience 2011–06–08, 31, doi:10.1523/JNEUROSCI.6273-10.2011.

26. Dutta Banik, D.; Martin, L.E.; Freichel, M.; Torregrossa, A.-M.; Medler, K.F. TRPM4 and TRPM5 are both required for normal signaling in taste receptor cells. Proceedings of the National Academy of Sciences 2018, 115, E772–E781, doi:10.1073/pnas.1718802115.

27. Ruan, Z.; Haley, E.; Orozco, I.J.; Sabat, M.; Myers, R.; Roth, R.; Du, J.; Lü, W.; Ruan, Z.; Haley, E., et al. Structures of the TRPM5 channel elucidate mechanisms of activation and inhibition. Nature Structural & Molecular Biology 2021 28:*7* 2021**–**06–24, 28, doi:10.1038/s41594-021-00607-4.

28. Sclafani, A.; Ackroff, K. Fat preference deficits and experience-induced recovery in global taste-deficient Trpm5 and Calhm1 knockout mice. Physiology & Behavior 2022/03/15, 246, doi:10.1016/j.physbeh.2022.113695.

29. Sclafani, A.; Zukerman, S.; Ackroff, K. Residual Glucose Taste in T1R3 Knockout but not TRPM5 Knockout Mice. Physiology & Behavior 2020/08/01, 222, doi:10.1016/j.physbeh.2020.112945.

30. Larsson, M.H.; Håkansson, P.; Jansen, F.P.; Magnell, K.; Brodin, P. Ablation of TRPM5 in Mice Results in Reduced Body Weight Gain and Improved Glucose Tolerance and Protects from Excessive Consumption of Sweet Palatable Food when Fed High Caloric Diets. PLOS ONE Sep 23, 2015, 10, doi:10.1371/journal.pone.0138373.

31. Richter, P.; Andersen, G.; Kahlenberg, K.; Mueller, A.U.; Pirkwieser, P.; Boger, V.; Somoza, V. Sodium-Permeable Ion Channels TRPM4 and TRPM5 are Functional in Human Gastric Parietal Cells in Culture and Modulate the Cellular Response to Bitter-Tasting Food Constituents. Journal of Agricultural and Food Chemistry February 20, 2024, 72, doi:10.1021/acs.jafc.3c09085.

32. Chubanov, V.; Köttgen, M.; Touyz, R.M.; Gudermann, T. TRPM channels in health and disease. Nature Reviews Nephrology 2024, 20, 175–187, doi:10.1038/s41581-023-00777-y.

33. Liu, Y.; Lyu, Y.; Wang, H. TRP channels as molecular targets to relieve endocrine-related diseases. 2024/01/01, 10.1016/B978-0-443-18653-0.00015-0, doi:10.1016/B978-0-443-18653-0.00015-0.

34. Jia, J.; Verma, S.; Nakayama, S.; Quillinan, N.; Grafe, M.R.; Hurn, P.D.; Herson, P.S.; Jia Jia, S.V., Shin Nakayama, Nidia Quillinan, Marjorie R Grafe, Patricia D Hurn, Paco S Herson. Sex Differences in Neuroprotection Provided by Inhibition of TRPM2 Channels following Experimental Stroke. Journal of Cerebral Blood Flow & Metabolism 2011–11, 31, doi:10.1038/jcbfm.2011.77.

35. Morales-Lázaro, S.L.; Rosenbaum, T. Multiple Mechanisms of Regulation of Transient Receptor Potential Ion Channels by Cholesterol. Sterol Regulation of Ion Channels 2017/01/01, 80, doi:10.1016/bs.ctm.2017.05.007.

36. Méndez-Reséndiz, K.A.; Enciso-Pablo, Ó.; González-Ramírez, R.; Juárez-Contreras, R.; Rosenbaum, T.; Morales-Lázaro, S.L.; Méndez-Reséndiz, K.A.; Enciso-Pablo, Ó.; González-Ramírez, R.; Juárez-Contreras, R., et al. Steroids and TRP Channels: A Close Relationship. International Journal of Molecular Sciences 2020*, Vol.* 21, Page 3819 2020–05–27, 21, doi:10.3390/ijms21113819.

37. Lakk, M.; Hoffmann, G.F.; Gorusupudi, A.; Enyong, E.; Lin, A.; Bernstein, P.S.; Toft-Bertelsen, T.; MacAulay, N.; Elliott, M.H.; Križaj, D. Membrane cholesterol regulates TRPV4 function, cytoskeletal expression, and the cellular response to tension. Journal of Lipid Research 2021/01/01, 62, doi:10.1016/j.jlr.2021.100145.

38. Lu, Y.-C.; Chen, C.-W.; Wang, S.-Y.; Wu, F.-S. 17β-Estradiol Mediates the Sex Difference in Capsaicin-Induced Nociception in Rats. The Journal of Pharmacology and Experimental Therapeutics 2009/12/01, 331, doi:10.1124/jpet.109.158402.

39. Ortíz-Rentería, M.; Juárez-Contreras, R.; González-Ramírez, R.; Islas, L.D.; Sierra-Ramírez, F.; Llorente, I.; Simon, S.A.; Hiriart, M.; Rosenbaum, T.; Morales-Lázaro, S.L., et al. TRPV1 channels and the progesterone receptor Sig-1R interact to regulate pain. Proceedings of the National Academy of Sciences 2018–2–13, 115, doi:10.1073/pnas.1715972115.

40. Carvalho, A.L.; Treyball, A.; Brooks, D.J.; Costa, S.; Neilson, R.J.; Reagan, M.R.; Bouxsein, M.L.; Motyl, K.J. TRPM8 modulates temperature regulation in a sex-dependent manner without affecting cold-induced bone loss. PLOS ONE Jun 4, 2021, 16, doi:10.1371/journal.pone.0231060.

41. Eckstein, E.; Pyrski, M.; Pinto, S.; Freichel, M.; Vennekens, R.; Zufall, F. Cyclic regulation of Trpm4 expression in female vomeronasal neurons driven by ovarian sex hormones. Molecular and Cellular Neuroscience 2020/06/01, 105, doi:10.1016/j.mcn.2020.103495.

42. Lin, F.; Liu, Y.; Rudeski-Rohr, T.; Dahir, N.; Calder, A.; Gilbertson, T.A.; Lin, F.; Liu, Y.; Rudeski-Rohr, T.; Dahir, N., et al. Adiponectin Enhances Fatty Acid Signaling in Human Taste Cells by Increasing Surface Expression of CD36. International Journal of Molecular Sciences 2023, *Vol.* 24, Page 5801 2023–03–18, 24, doi:10.3390/ijms24065801.

43. Dahir, N.S.; Calder, A.N.; McKinley, B.J.; Liu, Y.; Gilbertson, T.A. Sex differences in fat taste responsiveness are modulated by estradiol. Am J Physiol Endocrinol Metab 2021, 320, E566–E580, doi:10.1152/ajpendo.00331.2020.

44. Walmsley, R.; Chong, L.; Hii, M.W.; Brown, R.M.; Sumithran, P.; Walmsley, R.; Chong, L.; Hii, M.W.; Brown, R.M.; Sumithran, P. The effect of bariatric surgery on the expression of gastrointestinal taste receptors: A systematic review. Reviews in Endocrine and Metabolic Disorders 2024 25:*2* 2024**–**01–11, 25, doi:10.1007/s11154-023-09865-7.

45. Zhang, Y.; Hoon, M.A.; Chandrashekar, J.; Mueller, K.L.; Cook, B.; Wu, D.; Zuker, C.S.; Ryba, N.J.P. Coding of Sweet, Bitter, and Umami Tastes: Different Receptor Cells Sharing Similar Signaling Pathways. Cell 2003/02/07, 112, doi:10.1016/S0092-8674(03)00071-0.

46. Gilbertson, T.A. Gustatory mechanisms for the detection of fat. Current Opinion in Neurobiology 1998/08/01, 8, doi:10.1016/S0959-4388(98)80030-5.

47. Gilbertson, T.A.; Liu, L.; Kim, I.; Burks, C.A.; Hansen, D.R. Fatty acid responses in taste cells from obesity-prone and -resistant rats. Physiology & Behavior 2005/12/15, 86, doi:10.1016/j.physbeh.2005.08.057.

48. Gilbertson, T.A.; Fontenot, D.T.; Liu, L.; Zhang, H.; Monroe, W.T. Fatty acid modulation of K+ channels in taste receptor cells: gustatory cues for dietary fat. American Journal of Physiology-Cell Physiology 1997 Apr 01, 272, doi:10.1152/ajpcell.1997.272.4.C1203.

49. NS, D.; AN, C.; BJ, M.; Y, L.; TA, G. Sex differences in fat taste responsiveness are modulated by estradiol - PubMed. American journal of physiology. Endocrinology and metabolism 03/01/2021, 320, doi:10.1152/ajpendo.00331.2020.

50. Kim, J.W.; Roberts, C.; Maruyama, Y.; Berg, S.; Roper, S.; Chaudhari, N. Faithful expression of GFP from the PLCbeta2 promoter in a functional class of taste receptor cells. Chem Senses 2006, 31, 213–219, doi:10.1093/chemse/bjj021.

51. Zysow, B.R.; Kauser, K.; Lawn, R.M.; Rubanyi, G.M. Effects of Estrus Cycle, Ovariectomy, and Treatment with Estrogen, Tamoxifen, and Progesterone on Apolipoprotein(a) Gene Expression in Transgenic Mice. *Arteriosclerosis*, Thrombosis, and Vascular Biology 1997–09, 17, doi:10.1161/01.ATV.17.9.1741.

52. Nilsson, M.E.; Vandenput, L.; Tivesten, Å.; Norlén, A.-K.; Lagerquist, M.K.; Windahl, S.H.; Börjesson, A.E.; Farman, H.H.; Poutanen, M.; Benrick, A., et al. Measurement of a Comprehensive Sex Steroid Profile in Rodent Serum by High-Sensitive Gas Chromatography-Tandem Mass Spectrometry. Endocrinology 2015/07/01, 156, doi:10.1210/en.2014-1890.

