## Supplement Figure S1 for "Estrogen Dependent Variation in the Contributions of TRPM4 and TRPM5 to Fat Taste"

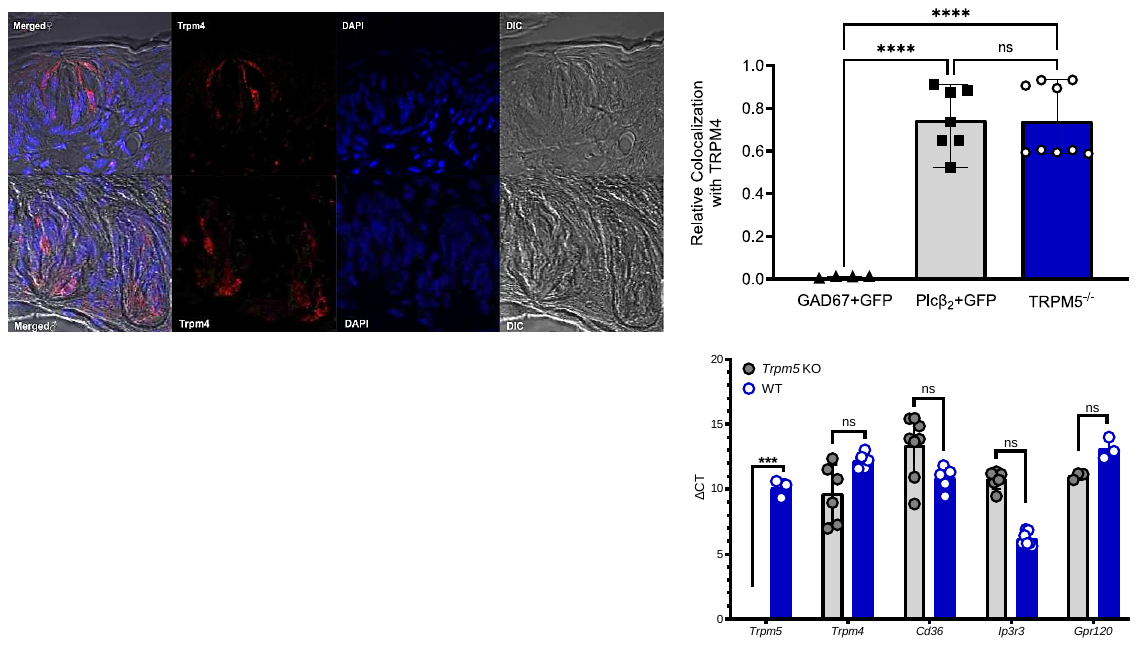

**Figure S1:** **TRPM4, IP_3_R3 are present in *Trpm5^-/-^* mice. (A)** Immunohistology images from male and female knockout mice indicate the presence of TRPM4 (upper panels) and IP_3_R3, shown for male mice (lower panels). Image groups denote merged, primary antibody (TRPM4 or IP_3_R3), DAPI, and differential contrast images (DIC) as collected using Nikon confocal microscopy. **(B)** Manders' colocalization coefficients for TBCs in GAD67 (Type III cells), Plcβ_2_ (Type II cells), and *Tprm5^-/-^* mice from NIS-elements analysis of images. Relative correlation coefficients are calculated by defining Regions of Interest (ROIs) that are carefully delineated around target structures, such as taste buds, and by establishing standard intensity thresholds to minimize background noise. Subsequently, the software calculates the relevant colocalization coefficients within the designated ROIs. Mander’s coefficients are calculated by summing the intensities of pixels in one channel only, where the intensity of the other channel is above zero, and then dividing by the total intensity of the respective channel. Mander’s coefficients allow for an assessment of how much of each protein or structure is found in the presence of the other target. These coefficients provide a robust, quantitative assessment of the degree to which each target protein or structure is found in the presence of the other. **(C)** Delta C_T_ (cycle threshold) values measured from qRT-PCR utilizing knockout and wild-type mice, asterisks denote no significant differences between WT and KO individuals from 2-way ANOVA with Tukey's post-hoc multiple comparisons. As expected, *Trpm5* was not detectable in taste cells from *Trpm5^-/-^* mice. Asterisks denote significant differences in ordinary one-way ANOVA with multiple comparisons: ***p<0.001, ****p<0.0001.
